# Maximum entropy methods for extracting the learned features of deep neural networks

**DOI:** 10.1101/105957

**Authors:** Alex Finnegan, Jun S. Song

## Abstract

New architectures of multilayer artificial neural networks and new methods for training them are rapidly revolutionizing the application of machine learning in diverse fields, including business, social science, physical sciences, and biology. Interpreting deep neural networks, however, currently remains elusive, and a critical challenge lies in understanding which meaningful features a network is actually learning. We present a general method for interpreting deep neural networks and extracting network-learned features from input data. We describe our algorithm in the context of biological sequence analysis. Our approach, based on ideas from statistical physics, samples from the maximum entropy distribution over possible sequences, anchored at an input sequence and subject to constraints implied by the empirical function learned by a network. Using our framework, we demonstrate that local transcription factor binding motifs can be identified from a network trained on ChIP-seq data and that nucleosome positioning signals are indeed learned by a network trained on chemical cleavage nucleosome maps. Imposing a further constraint on the maximum entropy distribution also allows us to probe whether a network is learning global sequence features, such as the high GC content in nucleosome-rich regions. This work thus provides valuable mathematical tools for interpreting and extracting learned features from feed-forward neural networks.

## Introduction

Multilayer artificial neural networks (ANNs) are becoming increasingly important tools for predicting outcomes from complex patterns in images and diverse scientific data including biological sequences. Recent works have applied multilayer networks, also called deep learning models, to predicting transcription factor (TF) binding [1, 2] and chromatin states [3–5] from DNA sequence, greatly advancing the state-of-the-art prediction rate in these fields. The success of these multilayer networks stems from their ability to learn complex, non-linear prediction functions over the set of input sequences. The main challenge in using deep learning currently resides in the fact that the complexity of the learned function coupled with the typically large dimension of the input and parameter spaces makes it difficult to decipher which input features a network is actually learning.

In spite of this challenge, the rise of deep learning has spurred efforts to interpret network prediction. While interpretation methods have been proposed in the fields of computer vision and natural language processing, we focus on genomics and specifically on methods for identifying features in individual input sequences used by an ANN for classification. To the best of our knowledge, two classes of interpretation methods have been proposed to address this problem: the first class of interpretation methods measures network feature importance by expressing the change in the network output for two distinct inputs as a sum of importance values assigned to the input units that encode biological sequence. One such decomposition assigns to each input unit importance given by its term in the 1^st^ order Taylor approximation of the change in the network output when the input units encoding a specific sequence are set to zeros. This method of assigning network importance to DNA sequence is called a Saliency Map by Lanchantin *et al*. [6] who adapted the method from the computer vision Saliency Map [7]. The DeepLIFT interpretation method uses an alternative approach for assigning input sequence importance based on comparing network activations elicited by the input sequences to those of a “reference” network input [8]. This approach to assigning importance has been motived as an approximation to Shapley values, which describe distribution of credit in game theory [9]. The second class of interpretation method, called *in silico* mutagenesis (ISM), measures changes in network output produced by simulated point mutations. Flexibility in the types of mutations performed means that ISM can, in principle, reveal the network’s dependence on sequence in detail. However, computational cost limits the number and type of progressive mutations that can be performed. As a result, to investigate learned network features using ISM, one must employ prior notions of important features to design a manageable number of sequential mutations for testing.

In this paper, we use the rigorous formalism of statistical physics to develop a novel method for extracting and interpreting network-learned sequence features. The method makes direct reference to the nonlinear function learned by the network by sampling a maximum entropy distribution over all possible sequences, anchored at an input sequence and subject to constraints implied by the learned function and by the background nucleotide content of the genome from which the network’s input sequences are derived.

To extract learned features from inputs, we study two complementary quantities derived from sequences sampled from the constrained maximum entropy (MaxEnt) distribution via Markov Chain Monte Carlo (MCMC): (1) a local profile of nucleotide contents for revealing sequence motifs, and (2) a feature importance score based on the sample variance of a summary statistic for focusing on a particular sequence characteristic of interest. The latter score directly measures the effect of a global sequence feature, such as GC content, on the non-linear function learned by the network, and it can be used to rank global features, thereby answering questions about the relative importance of such features in the context of network prediction.

Our approach can be viewed as a compromise between the two classes of interpretation method described above, extracting features by examining several sequences, instead of just two as in the first class, and eliminating ISM’s need for *a priori* specification of sequential mutations. Importantly, our method is distinguished from other previous approaches in that, rather than assigning an importance score to each base of a given input sequence, our method reveals features by sampling unseen sequences that are assessed by the trained network to be similar to the original input.

We apply our method, termed MaxEnt interpretation, to three separate deep neural networks and compare with the first class of interpretation methods: DeepLIFT and Saliency Map. The first network is a simple yet instructive example, and it demonstrates how the DeepLIFT and Saliency Map methods can encounter difficulties, while the MaxEnt method successfully captures the logic learned by the network. The remaining two networks are trained to predict transcription factor (TF) binding activity from CTCF ChIP-seq [2] and nucleosome positioning from chemical cleavage nucleosome mapping data [10], thus putatively learning the CTCF binding motifs and nucleosome positioning signals encoded in DNA, respectively. In the motif discovery task, while all methods give good results, correctly localizing the learned CTCF motifs in most cases, our MaxEnt approach achieves better agreement with motif positions called by the conventional motif discovery programs MEME and FIMO. In the task of extracting nucleosome positioning signals, MaxEnt surpasses DeepLIFT and Saliency Mapping in detecting a learned network preference for G/C and A/T nucleotides at positions separated by 10 base pairs (bps). Furthermore, we demonstrate the use of a global sequence feature score, unique to our method, to estimate the fraction of nucleosomal sequences for which the GC content is an important feature for network classification. We do not compare with ISM because, as discussed above, it is less general than other methods and requires the specification of the types of mutations to test.

## Results

### Constructing an input-specific constrained maximum entropy distribution

Consider a trained, multilayer feed-forward neural network that takes a length L genomic sequence as input and performs a classification task, assigning the sequence to one of *K* classes indexed by {0,1,…,*K*-1}. The network consists of a stack of layers, each of which contains a real-valued vector whose entries are called the activations of the layer’s units. The stack starts with an input layer whose activations represent the genomic sequence; activations of each subsequent layer are determined by applying a trained non-linear transformation to the proceeding layer. Activations of units in the output layer encode the predicted probabilities of the *K* classes, thereby assigning the input sequence to the class whose output unit activation is the largest. (S1 Text describes the types of layers used in this work)

The standard motivation behind multilayer networks is that intermediate layers may learn to recognize a hierarchy of features present in the set of inputs with features becoming more abstract with the depth of the intermediate layer. Since changes in the features detected by a layer are encoded in changes in the intermediate layer’s vector of activations, we propose that it is possible to identify learned features by looking for commonalities among the set of all input sequences that approximately preserve that layer’s vector of activations.

While this approach could be applied to identifying learned features of any intermediate layer, this work focuses on features learned by the penultimate layer, the layer making direct connections to the output layer. Penultimate layer features are interesting for two reasons. First, we are interested only in input sequence patterns that elicit an intermediate layer representation relevant to the classification task and discard sequence variations irrelevant to classification. Since the non-linear functions computed by intermediate layers are, in general, many-to-one mappings, one important role of a layer is to identify which differences in the preceding layer’s activations are irrelevant to the learned classification. Because intermediate layer activations are calculated from the input sequence by composing such non-linear functions, the number of identified classification-irrelevant differences in inputs should increase with the intermediate layer depth, making the penultimate layer the intermediate layer least affected by classification-irrelevant features. Second, the network output layer is either a logistic or a softmax classifier applied to penultimate activations; by uncovering features learned by the penultimate layer, we are thus finding the sequence patterns directly used by these output layer classifiers to make predictions.

To formalize the search for new input sequences that approximately preserve a given penultimate activation, let the vector ***x***_**0**_ represent an input genomic sequence. The entries in ***x***_**0**_ encode the presence of one of the nucleotides A, C, G, T at a certain position in the sequence. Let ****Φ**** ***x***_**0**_ denote the vector of penultimate layer activations elicited by ***x***_**0**_. For notational convenience, assume the trained network has assigned ***x***_**0**_ to the 0^th^ class. We measure the extent to which an arbitrary input sequence ***x*** reproduces the penultimate activation ***Φ x***_**0**_ by weighing the set of all length *L* genomic sequences with the probability mass function (PMF) 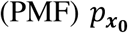 that is most similar to a pattern-agnostic PMF *q*, subject to a constraint on the average distance to ***Φ x***_**0**_. More precisely, we define

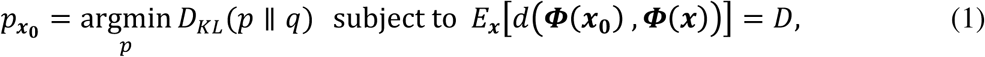

where *D*_KL_(*p* ∥ *q*) is the Kullback-Leibler (KL) divergence between *p* and *q*, *E*_***x***_ denotes expectation calculated when ***x*** is distributed according to *p*, *d* is a distance metric on the set of penultimate activation vectors, and *D* is a positive constant, with smaller values corresponding to a more exact reproduction of the activation vector ***Φ x***_**0**_. The PMF *q* describes prior beliefs about the background nucleotide content of the genomic regions from which network inputs are derived, and the fact the 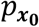 minimizes *D_KL_*(*p* ∥ *q*) subject to the constraint ensures that differences between 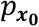 and these prior beliefs arise from the need to reproduce the sequence features encoded in ***Φ x***_**0**_. We take *q* be a product of *L* identical single nucleotide distributions with probabilities of G/C and A/T chosen to reflect the genome-wide GC content. In this case, the solution to (1) is

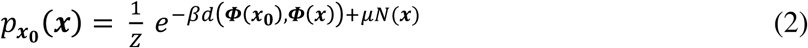

where *Z* is a normalization constant, *β* is a Lagrange multiplier whose value is chosen to yield the desired value of *D*, *N*(***x***) is the number of G and C nucleotides in sequence ***x***, and 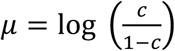 where *c* is the GC content of the genome (S2 Text). When *μ* = 0, 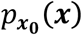 ***x*** is determined solely by the learned function ***Φ***, and equation (2) is the maximum entropy distribution over all length *L* sequences subject to the constraint in (1). For simplicity, we use the term MaxEnt samples to refer to the samples from 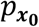 for any value of *μ*.

We use a weighted Euclidean metric for *d*,

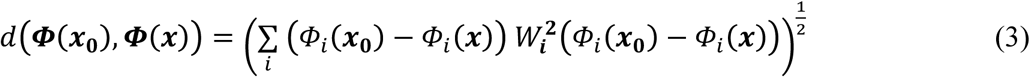

where our choice of *W_i_* depends on the type of classification task. For binary classification, *W_i_* = *w*_0,*i*_, the weight connecting the *i*^th^ penultimate unit to the output unit encoding the assigned class label of ***x***_**0**_ (when there is only one output unit, *W*_d_ is the weight of connection to this unit). For multiclass classification, *W_i_* = *w_0,i_* – *w_k,i_*, where *k* ∈ {1,2, … *K* – 1} is a user-specified class. This choice of *W*_d_ corresponds to weighting each penultimate activation by its effect on the log ratio of predicted class 0 and class *k* probabilities (S1 Text).

The mean distance *D* is an increasing function of 1/*β*, whose scale is set by nearest-neighbor distances among penultimate activations, as measured by the metric (3). When *β* is large, *D* approaches 0, the PMF over the set of penultimate activations is a single spike at ***Φ x***_**0**_, and 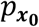 consists of a (relatively) small number of non-zero probability masses on sequences ***x*** that exactly reproduce ***Φ x***_**0**_. Conversely, decreasing *β* smooths the PMF over penultimate activations and causes 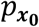 to shift probability mass onto sequences that yield penultimate activations similar to ***Φ x***_**0**_. When 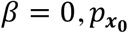 and *q* are identical, *D* is the expected distance under the distribution *q*, and 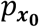 contains no information on the features encoded in ***Φ x***_**0**_.

This intuition informs one method for choosing *β* (and implicitly *D*): select *β* so that the PMF over the set of penultimate activations has small width relative to an empirical distribution of penultimate activations, while still assigning appreciable probability to sequences with penultimate activations near ***Φ x***_**0**_. Alternatively, because a sufficiently large value of *β* effectively fixes the nucleotide content at certain indices in sequences sampled from 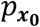, one can examine the samples from distributions at different values of *β* to uncover a hierarchy of important features in ***x***_***0***_. We give examples of both methods in the following sections, where we sample the distribution (2) using MCMC (Methods). Figure 1, illustrates the sampling of sequences ***x*** according to their similarity to data set element ***x***_**0**_ in the space of penultimate layer activations.

**Figure 1.**
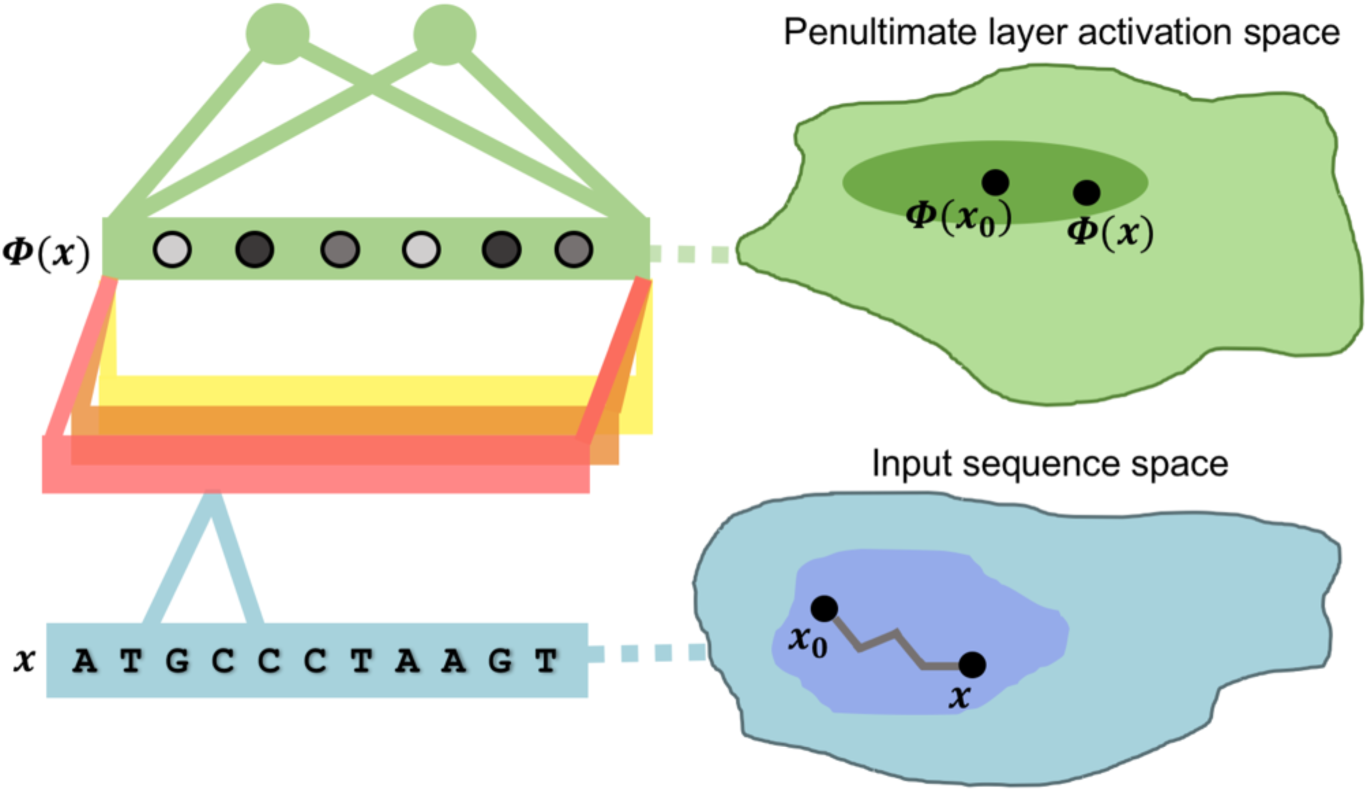
Schematic representation of our MaxEnt interpretation method.

An unseen sequence ***x*** elicits penultimate unit activations (shaded dots in left figure) via non-linear operations of intermediate layers (illustrated as a horizontal stack of convolutional filters). The MaxEnt method for interpreting a given input sequence ***x***_**0**_ assigns probability to a new sequence ***x*** according to its similarity to ***x***_**0**_ in the space of penultimate activations. The irregular path connecting ***x***_**0**_ and ***x*** in sequence space illustrates the steps of MCMC.

### Extracting features from samples

We extract features of network input ***x***_**0**_, captured by penultimate activation ***Φ***(***x***_**0**_), by examining several statistics of the MCMC samples from 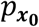. In our first example, the dimension of the input space is low enough to directly visualize the empirical distribution of samples. For higher dimensional input spaces, we summarize the distribution by plotting sample single nucleotide frequencies at each genomic index and also by examining the variance of linear combinations of nucleotide indicator variables that serve to define sequence features of interest.

Plots of single nucleotide frequencies reveal important features by illustrating the extent to which preserving penultimate activation forces the single nucleotide distribution of 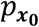 to be away from that of *q.* Large divergence of sample nucleotide frequencies from *q* signals importance and determines which nucleotide substitutions dramatically affect the penultimate layer representation. By contrast, if the sampled nucleotide distribution of 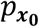 is similar to that of *q* at a given base position, and if we assume that the content of the position is independent of other positions under 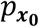, then this position is irrelevant for determining ***Φ***(***x***_**0**_) and the classification of ***x***_**0**_.

To quantify the importance of sequence features, in a way that accounts for interactions among base positions, we define an input-wide sequence feature *V* as a function that maps an input sequence to a weighed sum of indicator variables for specific nucleotide content:

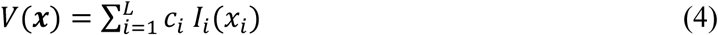

where *x_i_* denotes the nucleotide at index *i* in ***x***, *I_i_*(·), is the indicator variable for one of a set of nucleotides at index *i* and *c_i_* is a real valued weight. We define the concordance of sequence ***x*** with the input-wide feature *V* to be *V*(***x***). For example, if we are interested in the importance of GC content, *V* would be the sum of indicator variables for S (G or C nucleotide) at each input index, and ***x*** would have large concordance with this input-wide sequence feature when it contains many G and C nucleotides.

To relate changes in feature concordance to changes in penultimate layer activation, let *X*_v_ denote the set of length *L* sequences ***x***, such that *V* (***x)*** = *v*. The quantity

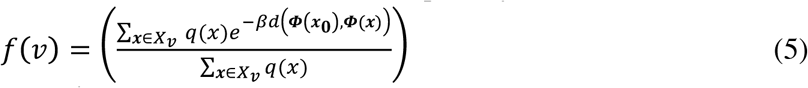

is the mean value of 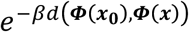 under PMF *q*, conditioned on *V*(***x)*** = *v*. The rate of decay of *f* (*v)* from its maximum measures the dependence of *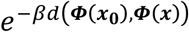 V*(***x***). To see this, observe that when *V* sums indicator variables for nucleotides important in eliciting the penultimate activation ***Φ***(***x***_**0**_), the exponential factor *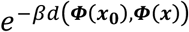* ^***x***^ will, on average, decay as *V*(***x***) deviates from *V*(***x***_**0**_). More rapid decay corresponds to greater importance, since in this case smaller changes in *V*(***x***) suffice to produce large changes in ***Φ* (*x*)**. By contrast, when *V* sums indicator variables for only those nucleotides unimportant in eliciting ***Φ***(***x***_**0**_), the factor *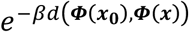* does not decay with changes in *V* (***x)***.

In S2 Text, we approximate the decay of *f* from its maximum as

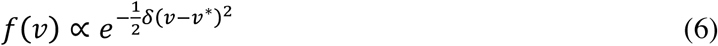

where *v*^*∗*^ maximizes *f*(*v*) and

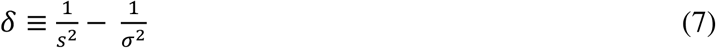

where *s*^2^ is the variance of *V*(***x***) under the PMF *p*_***x***_ and *σ*^2^ is the variance of *V*(***x***) under PMF *q*. The variance *s*^2^ is estimated from MCMC samples while the variance *σ*^2^ can be calculated explicitly from *q*. Larger values of *δ* correspond to more rapid decay of *f v*, signaling greater input-wide feature importance.

Our derivation of the proportionality (6) requires that the marginal distributions of *V*(***x***) when ***x*** is distributed according to *q* or 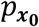 are approximately normal. Approximate normality of *V*(***x***) when ***x*** is distributed according to *q* is guaranteed by the Lindeberg version of the Central Limit Theorem, provided that *V* sums indicator variables at a large number of base positions with weights roughly equal in magnitude. Approximate normality of *V*(***x***) when ***x*** is distributed according to 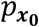 can be checked directly by estimating *V*(***x***) from MCMC samples. We have checked that these normal approximations are valid when the number of base positions considered by the function *V* is large, explaining our choice of the name input-wide sequence features.

In Application 3 below, we consider two uses of the importance measure *δ*. First, we choose the weights *c_i_* and indicators *I_i_* (·), so that the resulting input-wide feature *V* measures the importance of GC content for a network predicting nucleosome positioning. Second, we use MCMC samples from (2) to find sets of weights *c*_d_ that approximately maximize *δ*. When *V* captures an important input-wide feature, the exponential factor *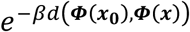*in (2) should make the marginal distribution of *V* under 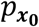 much narrower than the marginal distribution of *V* under *q*; that is, *s*^2^*≪ σ*^2^. In this case, we can thus approximate,

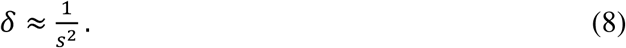

Under this approximation, the most important features are given by the lowest variance principal components (PCs) of our MCMC samples. Examining the elements of these low variance PC vectors reveals important input-wide features.

### Application 1: Interpreting learned XOR logic

ANN interpretation methods, such as Saliency Map and DeepLIFT, that assign a real-valued importance score to each input unit provide intuitive pictures that are easy to understand, but must confront the challenge of summarizing learned nonlinear interactions between base positions using a base-wise score. To illustrate the practical consequences of these issues, we trained ANNs on an artificial data set of DNA dinucleotides and applied the MaxEnt, Saliency Map and DeepLIFT interpretation methods. Class labels were assigned to each of the 16 possible dinucleotides according to an XOR logic, where sequences (with positions indexed by 0 and 1) were assigned to class 0 unless one, but not both, of the following conditions was satisfied, in which case class 1 was assigned:

- sequence position 0 is W (A or T),
- sequence position 1 is G.

We represented sequences with a one-hot encoding scheme and chose a simple convolutional architecture with 2 convolutional filters of stride 1 and taking a single base of the dinucleotide as input, followed by a layer of two fully connected units and then a single output unit indicating the predicted class label. Rectified-linear units (ReLU) were used throughout the network, except for the output which was modeled using a sigmoid function (S1 Text). We obtained 30 of these convolutional architectures trained to achieve 100% classification accuracy on the set of all possible dinucleotide inputs (Methods).

Figure 2 shows the results of interpretation analysis on the AA and GG inputs in class 1 for each of the 30 models achieving 100% validation accuracy. Although each network represents the same classification rule, the interpretation results of Saliency Map and DeepLIFT show model dependence in the sign of their interpretation score, indicating in some cases that a nucleotide is evidence for the correct class label and in other cases that the same nucleotide is evidence against this class label (Figure 2A-D) (see S3 Text for description of our application of Saliency Map and DeepLIFT).

**Figure 2.**
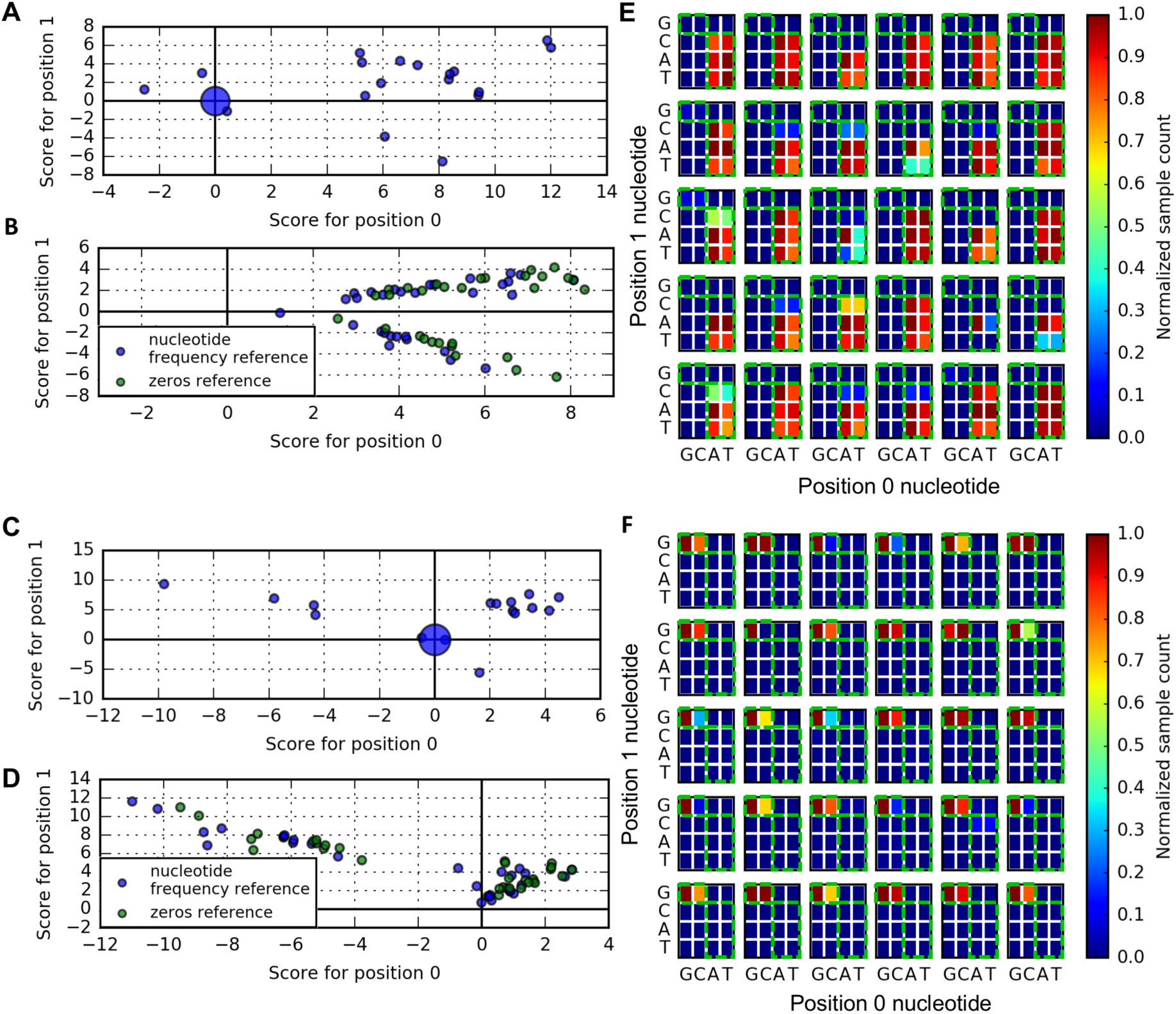
Interpretation of XOR network inputs. (A, B) Scatter plots of interpretation scores assigned to the 0^th^ and 1^st^ sequence position by Saliency Map and DeepLIFT interpretation, respectively, for the AA network input. Markers at the origin have size proportional to number of overlapping data points. Colors in **(B)** indicate DeepLIFT interpretation scores using different reference inputs (see S3 Text). **(C, D)** Same as **(A, B)**, respectively, but for the GG network input. **(E)** Density of MCMC samples from MaxEnt distribution (2) for AA input. Densities are normalized by the most abundant dinucleotide. Green boxes highlight the set of dinucleotide inputs belonging to class 1. **(F)** Same as **(E)** but for the GG network input. All results are interpretation of the same 30 ANNs.

In contrast, our MaxEnt interpretation illuminates, in almost every case, how the trained networks implement the XOR logic defined above, indicating that the GG input is similar to CG and that the AA input is similar to any input with A or T in the 0^th^ position and not G in the 1^st^ position (Figure 2E,F). For this analysis, we sampled the distribution (2) with *μ* = 0 and *β* chosen based on the distribution of penultimate layer activation associated with the 16 dinucleotide inputs (Methods). Figure S1 shows similar results for the other dinucleotide inputs.

Figure 2 highlights a key difference that distinguishes the MaxEnt interpretation approach from Salience Map and DeepLIFT. By replacing base-position scores with samples from a distribution, MaxEnt interpretation is able to capture nonlinear classification rules that escape the other methods. The cost of this extra flexibility is some additional effort in assigning meaning to the MaxEnt samples.

### Application 2: Localizing learned motifs

We applied MaxEnt interpretation to a network trained on a benchmark motif discovery data set constructed by [2] from ENCODE CTCF ChIP-seq data [11]. CTCF is a well-studied DNA binding protein with important transcription factor and insulator functions [12, 13]. In this motif discovery task, the network distinguished elements of the positive class, consisting of 101 base-pair (bp) sequences centered on ChIP-seq peaks, from elements of the negative class consisting of positive class sequences shuffled to maintain dinucleotide frequency. We represented network inputs with a one-hot encoding scheme and trained an architecture consisting of a convolutional layer of 64 convolutional filters each with a stride of 1 and taking 24 bps as input, followed by a layer of 100 units fully connected to the preceding layer and a two unit softmax output layer. ReLUs were used in all layers preceding the output (S1 Text). The trained network performed well, achieving a mean area under the receiver operating characteristic (AUROC) of 0.978 with standard deviation 0.001 in 5-fold cross-validation (Methods).

We picked a neural network trained on a random fold of the cross-validation, selected 2500 random sequences from all correctly classified CTCF-containing sequences in the test set, and applied MaxEnt, DeepLIFT and Saliency Map interpretation methods. Our application of MaxEnt to this network used *β* = 400, chosen by examining samples collected at a range of *β*′*s* for a few network inputs and selecting the smallest *β* sufficient to fix the nucleotide content at positions where MaxEnt marginal distributions signaled greatest importance. Because single nucleotide frequencies for the data set were nearly uniform (P(A) = P(T) = 0.27 and P(C)=P(G) = 0.23), we set *μ* = 0 when sampling from the distribution (2). Figures 3A and 3B show nucleotide frequencies as a function of base index for MCMC MaxEnt samples associated with two input sequences. In both cases, the location of the motif identified by the network was indicated by an interval of single nucleotide frequencies that diverged dramatically from the uniform distribution over nucleotides implied by the distribution (2) for sequence locations with little effect on penultimate layer activations. Sequence logos were generated from the nucleotide frequencies on these intervals using WebLogo [14]. We confirmed that the discovered motifs in Figures 3A and 3B correspond to the canonical CTCF motif and its reverse complement by using the motif database querying tool Tomtom [15] (Methods, Figure S2).

**Figure 3.**
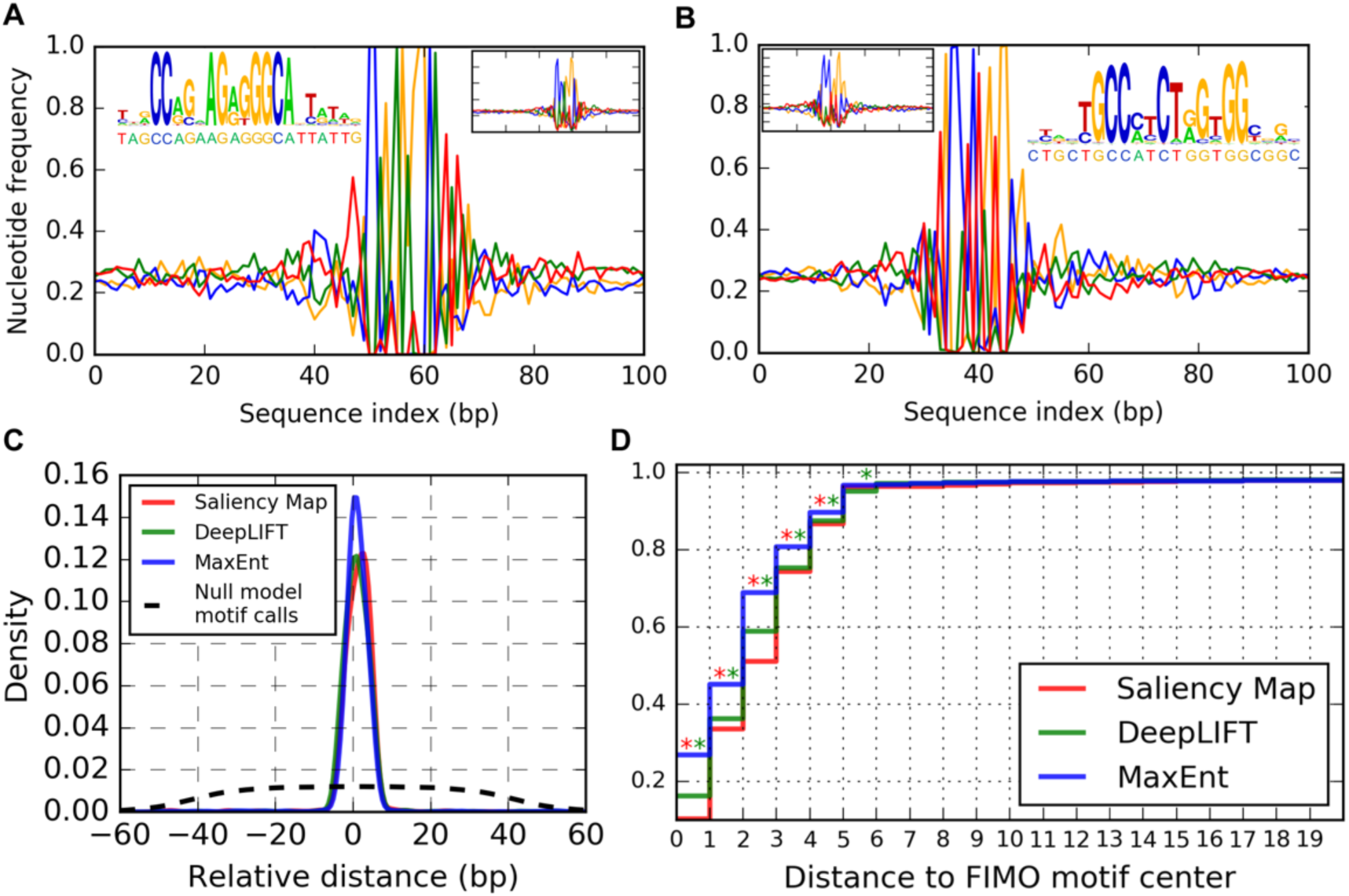
Interpretation of CTCF-bound sequences. (A,B) Nucleotide frequencies of MCMC samples from MaxEnt distribution (2) for two input sequences that the ANN correctly identified as CTCF bound. Main plots correspond to sampling at *β*= 400; inset line plots correspond to sampling at *β*= 100, illustrating the multiscale nature of our interpretation method. Inset sequence logos show the called motifs, with the corresponding input sequences indicated below the horizontal axis. Colors green, blue, orange, red correspond to A, C, G, T. **(C)** Kernel-density smoothed distribution of relative distances between motifs called by network interpretation methods and motifs called by FIMO. Null model density is estimated by calling motif positons with uniform probability over the set of 19bp intervals contained in the 101 bp network inputs. (**D)** Cumulative distribution of the absolute distances from **(C)**. Red asterisk at (x,x+1) indicates significantly fewer Saliency Map motif calls than MaxEnt motif calls within x bp from a FIMO motif (one-sided binominal test, p < 0.01)). Green asterisks indicate the similar comparison between DeepLIFT and MaxEnt motif calls.

DeepLIFT and Saliency Map interpretation of these inputs yielded visually similar results (Figure S3). However, MaxEnt single nucleotide frequencies provide direct interpretation as motif position-specific scoring matrices utilized by other bioinformatics tools and thus provide advantages over Saliency Map and DeepLIFT base-wise scores.

To make a more global comparison of interpretation methods, we calculated the distribution of relative distances from motif positions called using each interpretation method to motif positions identified by the conventional motif discovery programs MEME and FIMO [16, 17]. Combined application of MEME and FIMO to the 2500 interpreted inputs found a single 19 bp consensus CTCF motif in 1652 of these input sequences (Methods). Using this motif length as a guide, we called MaxEnt motifs by calculating, for each sequence input, the KL divergence of sample nucleotide frequencies at each base from a uniform distribution and finding the 19 bp running window that has the largest average KL divergence. DeepLIFT and Saliency Map motifs were similarly called for each sequence input at the 19 bp widow with largest interpretation score (see S3 Text for definition). Figure 3C shows the empirical distribution of the signed center-to-center distances between network interpretation motifs and MEME/FIMO motifs in the 1652 sequences. Figure 3D shows the cumulative distribution of the same unsigned distances. MaxEnt interpretation gives significantly more motif calls within 0, 1, 2, 3 and 4 bp of MEME/FIMO motifs than Saliency Map and DeepLIFT interpretation.

### Application 3: Extracting nucleosome positioning signals

Finally, we tested ANN interpretation methods on a genomic data set where we expected learned sequence features to be more diffuse. We constructed a data set based on the chemical cleavage map of nucleosome dyads in *S. cerevisiae* [10]. Each input was a 201 bp sequence with positive class elements centered on nucleosome dyads and negative class elements centered on locations uniformly sampled from genomic regions at least 3 bps from a dyad. We chose to allow sampling of negative sequences within the 73 bps of nucleosomal DNA flanking the dyad to encourage the network to learn features that direct precise nucleosome positioning as well as those determining nucleosome occupancy.

Our trained network consisted of a convolutional layer with 30 filters, each with a stride of 1 and taking 6 bp windows as input, followed by a 400-unit layer with full connections and a 2-unit output softmax layer. Sigmoid activation functions were used in all layers preceding the output (S1 Text). The trained network performed well, achieving an AUROC of 0.956 on the test set (Methods). We applied interpretation methods to 2500 input sequences randomly selected from validation set elements corresponding to nucleosomal sequences correctly classified by the network. MaxEnt interpretation sampled the distribution (2) using *μ* = –0.49 and *β* = 40.0. The value of *μ* was determined using the 38% GC content of the *S. cerevisiae* genome. The value of *β* was determined by examining the plots of nucleotide frequencies for a range of *β* values and selecting the largest value that permitted fluctuation in the nucleotide content at all of 201 bp. S3 Text describes our application of DeepLIFT and Saliency Mapping to this network.

Figure 4A shows single nucleotide frequencies of MaxEnt samples for one of the 2500 nucleosomal sequences analyzed. Figures 4B and 4C show the results of DeepLIFT and Saliency Map interpretation for the same network input, respectively. This example shows that MaxEnt interpretation surpasses DeepLIFT and Saliency Mapping in capturing the importance of G/C and A/T nucleotides preferentially positioned at anti-phased 10 bp intervals. To confirm this trend across all interpreted nucleosomal sequences, we calculated for each input the Discrete Fourier Transform (DFT) of single nucleotide frequencies of MaxEnt samples, DeepLIFT interpretation scores, and Saliency Map interpretation scores (S3 Text). DFT represents each of these signals as a sum of sinusoidal functions, with each sinusoid described by a period, phase and amplitude of oscillation. The contribution of a sinusoid to the signal is measured by its amplitude relative to the amplitudes at all periods; we normalized the components of each DFT to account for this relative comparison (Methods). Figure 4D shows the average of these normalized amplitudes over the set of all interpreted inputs, confirming that MaxEnt single nucleotide frequencies provide the strongest evidence for the learned 10bp-periodicity preference in nucleotide positioning.

**Figure 4.**
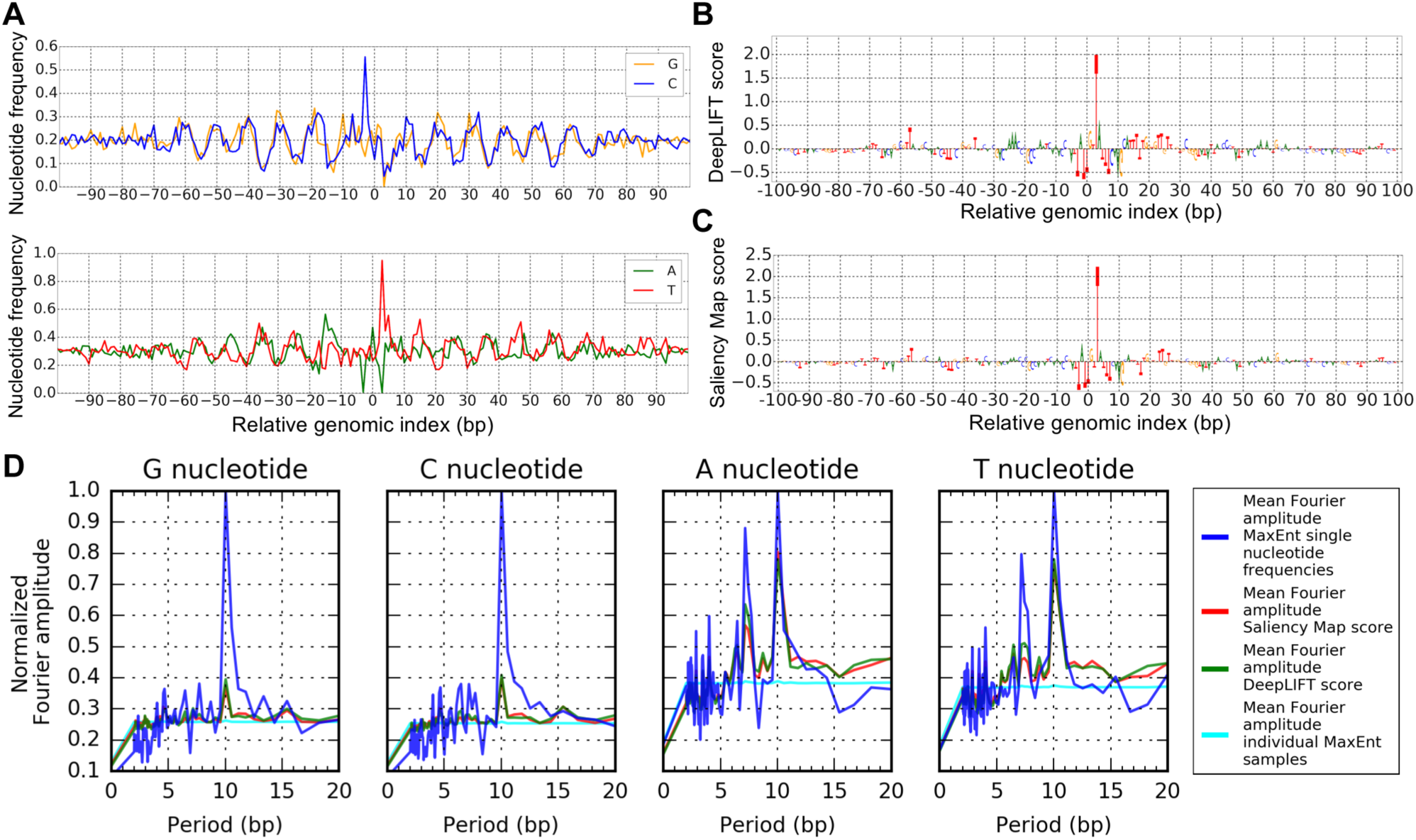
Interpretation of nucleosome positioning signals. (A) Nucleotide frequencies for samples from MaxEnt distribution (2) associated with a single nucleosomal input sequence. **(B)** DeepLIFT interpretation scores for the input analyzed in **(A). (C)** Saliency Map interpretation scores for the input analyzed in (**A**) (representation of DeepLIFT and Saliency Map scores uses code from [8]). **(D)** Normalized Fourier amplitudes of interpretation scores averaged over 2500 interpreted nucleosomal sequences correctly classified by the network. Note vertical axis is scale by maximum value.

Consistent with Figure 4D, 10 bp-periodic signals were found in many individual sets of MaxEnt samples. However, Figure S4 shows counterexamples to this trend, highlighting that the network does not need to detect the pattern to classify a sequence as nucleosomal. Figure S4 also shows a plot of nucleotide frequencies averaged over the set of 2500 input nucleosomal sequences.

It is important to recognize that even though these plots of sample nucleotide frequencies illuminate learned features, they do not imply that the periodic features are present in individual MaxEnt samples. Indeed, it has recently been shown that most nucleosomal sequences in *S. cerevisiae* do not contain significant 10 bp periodicity [18]. To confirm that the MaxEnt samples produced by our method were also not individually enriched for 10 bp periodicity, we calculated the normalized Fourier spectrum of each sample separately and then averaged the amplitudes over all samples associated with the 2500 nucleosomal sequences (Methods). Figure 4D shows that this Fourier amplitude at 10 bp averaged over the pooled MCMC samples is greatly suppressed relative to the Fourier amplitude of nucleotides frequencies averaged over nucleosomal sequences. In this way, MaxEnt samples capture the true nature of the 10 bp periodic feature learned by the network. That is, to be considered similar to an input nucleosomal sequence, it is enough for MaxEnt samples to possess G/C and A/T nucleotides at only *some* of the “hot-spots” separated by 10 bps; at the same time, averaging over these samples gives a coarse and conceptually useful representation of the learned feature.

It is also widely believed that nucleosomal DNA often possesses high GC content relative to the genomic background [19]; we thus explored the importance of GC content to our network’s classification. Figure 5A shows that, while mean GC content of MaxEnt samples generally agreed with the background 38% tuned by our choice of *μ*, there was also a significant positive correlation between sample mean GC content and GC content of the associated input. The correlation indicated that changes in GC content affected the penultimate layer activations to the extent that samples tended to preserve the GC enrichment or depletion of their associated input.

**Figure 5.**
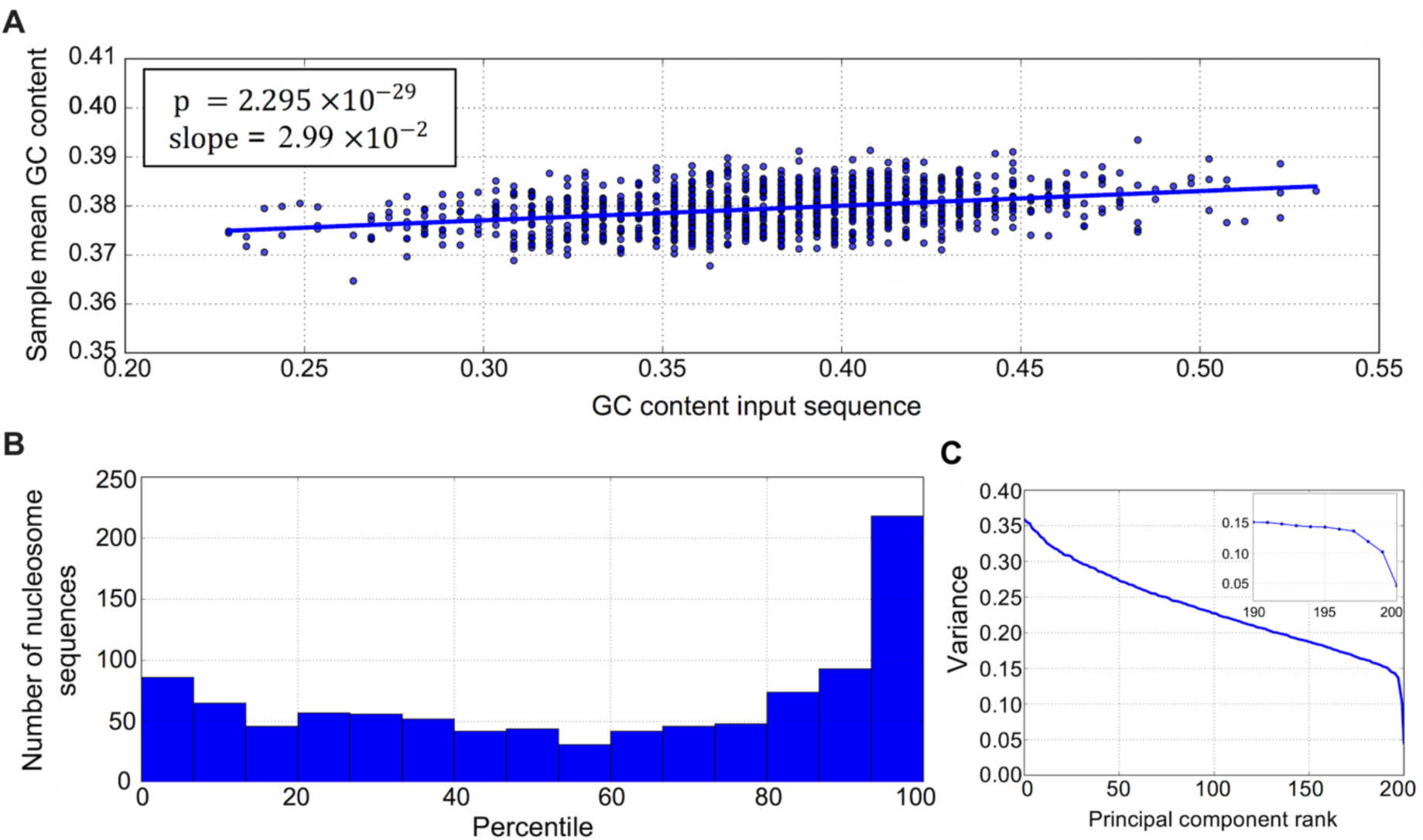
Assessing the importance of GC content for nucleosome positioning. (A) Distribution of GC content of 1000 network input sequences corresponding to nucleosomal DNA and the mean GC content of samples associated with these inputs. **(B)** Histogram of the percentiles of GC feature importance scores in the distribution of importance scores of 300 “dummy” sequence features. Histogram summarizes percentiles of GC feature importance scores for 1000 nucleosomal sequences. **(C)** Example of decay in variance associated with ranked principal component vectors in PCA analysis of samples from the distribution (2).

To rigorously measure the importance of the GC content feature, we defined an input-wide sequence feature *V* that sums the indicators for G or C nucleotides at each of the central 147 bases of the network input. For comparison, we defined “dummy” input-wide features which also sum indicator variables at each of the central 147 bases of network input, but where, at each position, the set of two nucleotides for which the indicator is 1 is uniformly sampled from the list {G,C}, {G,A}, {G,T}, {C,A}, {C,T}, {A,T}. For 1000 inputs chosen at random from the 2500 analyzed, we calculated the feature importance score *δ*, defined in (7), for the GC content feature *V* and for 300 random variables measuring dummy features. We then computed the percentile of the importance score of the GC content variable in the distribution of importance scores of the dummy feature variables for each input. Figure 5B shows the distribution of these percentiles, with enrichment of nucleosomal sequences near the 100^th^ percentile; setting a threshold at the 90^th^ percentile in the distribution of dummy feature importance scores, we estimate that GC content is a learned network feature of about 26% of the 1000 nucleosomal sequences analyzed.

While assigning relative importance to a chosen input-wide feature is useful, we were also interested in automatically discovering the most important input-wide features from MaxEnt samples, without prior specification of the weights *c_i_* in (4). For this purpose, we chose *V* to sum over indicator variables for G/C at each position, with the *c_i_*’s to be determined. The variance of this *V* ***x***, with ***x*** distributed according to (2), can be written for an arbitrary vector ***c*** ≡(*c*_1_, *c*_1_, …, *c*_*L*_)^*T*^of weights as

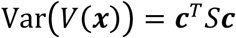

where *S* is the covariance matrix of the indicator variables estimated from MCMC samples. Since the approximation (8) implies that feature importance decreases with increasing variance, we seek weight vectors minimizing Var (*V(* ***x***)) under the constraint that ***c*** has unit Euclidean norm. Thus, the problem of identifying low-variance input-wide features amounts to selecting low variance principal components (PCs) in Principal Component Analysis (PCA) using *S*. The elements of the low variance PCs then give the weights of an input-wide feature *V*. Moreover, because the PCs are uncorrelated with respect to *S*, we expect several of the low variance PCs to be interpretable. Figure 5C shows a sharp decrease in the variance of the input-wide features determined by PCA on MaxEnt samples for a single nucleosomal sequence. We empirically observed that this sharp drop, signaling the prominent importance of features corresponding to the lowest variance PC vectors, is typical for our samples.

Figure S5 plots the weight vectors obtained from the two lowest variance PC vectors associated with the MaxEnt distribution depicted in Figure 4A. The lowest variance feature concentrates on a spike at the +3 position relative to the dyad. The strong network dependence on this position is also seen in Figure 4 A-C. The second lowest variance feature shows 10 bp periodicity with the weights changing sign roughly every 5 bp. While the pattern is much like that of Figure 4A, it accounts for correlations in nucleotide content, demonstrating that it is the collective alignment of G/C and A/T content with this weight template that changes the network’s penultimate representation.

Finally, we demonstrated the utility of these features by constructing a simple 10-nearest neighbor classifier, where we used the lowest variance PCs to compute the inter-sequence distance. Briefly, we randomly selected 1200 correctly classified nucleosomal sequences to which we applied our interpretation method with the values of *β* and *μ* given above. For a given nucleosomal sequence, we represented each of its MCMC samples as a 201 dimensional binary vector by evaluating the G/C indicator variable at each base and used the element-wise mean over these vectors to represent the nucleosomal sequence itself as a positive exemplar in the nearest neighbor classification. Likewise, we repeated this task for 1200 correctly classified non-nucleosomal sequences to obtain negative exemplars. We then selected a balanced test set of 10500 sequences that were previously classified correctly by the network and represented each test sequence as a 201 dimensional binary vector indicating the G/C nucleotide composition of its bases. To compute the distance of a test sequence to an exemplar, we projected the vector joining the exemplar and the test sequence onto the space spanned by the exemplar’s 5 lowest variance PC vectors, scaling the projected coordinates by the inverse standard deviation of the associated PC vectors and then computing the Euclidean distance. Test set elements were assigned to the majority class of the 10 nearest exemplars. This simple method yielded a classification accuracy of 76%. For comparison, we repeated this classification replacing the 5 lowest variance PC vectors of each exemplar with 5 mutually orthogonal vectors randomly sampled from the 201 dimensional space (Methods). Using this control, nearest neighbor classification accuracy dropped to 51%. This result thus demonstrates the ability of our interpretation method to extract *de novo* features used in the neural network’s classification.

## Discussion

Deep neural networks provide researchers with powerful tools for making predictions based on complex patterns in biological sequence. Methods for extracting learned input features from these networks can provide valuable scientific insights, and several efforts in this direction [6, 8] have made deep learning an even more appealing approach for tackling complex problems in genomics and other scientific disciplines.

We have contributed to these efforts by introducing a novel feature extraction method based on sampling a maximum entropy distribution with a constraint imposed by the empirical non-linear function learned by the network. From a theoretical standpoint, this constraint allows the derivation of relationships between the statistics of the sampled distribution and the dependence of network classification on specific sequence features. In particular, we have developed a scheme for assessing input-wide feature importance that has been difficult to measure otherwise with currently available approaches to network interpretation.

From a practical standpoint, the MaxEnt approach to feature extraction is distinct from other interpretation methods that assign base-wise importance to sequences. Admittedly, different interpretation schemes may thus have distinct advantages and disadvantages. For example, in Application 1, the MaxEnt method is able to capture the XOR logic that is learned by a simple ANN, but the same logic is difficult to infer using the methods based on base-wise importance. In Application 2, all schemes give similar results, but the MaxEnt interpretation method also provides probabilistic position-specific scoring matrices that are commonly used in bioinformatics. However, DeepLIFT and Saliency Map may be preferred in some cases for their computational efficiency. Interpreting an input in Application 2 via DeepLIFT, Saliency Map and MaxEnt takes 0.64 ms, 0.11 ms, and 181 s, respectively, on a quad-core 3.2 GHz Intel CPU, where the clear computational cost of MaxEnt interpretation stems from our MCMC sampling approach. This cost could be mitigated via a parallel implementation of multiple MCMC chains. Finally, Application 3 illustrates a setting in which MaxEnt interpretation surpasses other methods in elucidating the learned features that are consistent with the current understanding of nucleosome positioning [18].

The success of our MaxEnt approach signals that statistical physics may have much to contribute to the task of interpreting deep learning models. Indeed, a central goal of statistical mechanics is to understand constrained MaxEnt distributions of many degrees of freedom that interact according to known microscopic rules. While our formulation addresses an inverse problem of inferring unknown characteristics of network inputs from observed statistics of a constrained MaxEnt distribution, statistical mechanics provides a wide range of tools that could be further explored in this new context. This theoretical paper provides an example of the growing synergy between machine learning and physics towards assessing the role of diffuse and subtle sequence features that direct important biological outcomes, such as the positioning of nucleosomes.

## Methods

### Monte Carlo sampling

Given a representation ***x***_**0**_ of a sequence classified to class 0 by the trained network, we sampled sequences ***x*** from the PMF (2) using a Markov chain Monte Carlo (MCMC) method. We initialized the Markov random sequence ***x*** at ***x***_**0**_, then repeatedly selected an index *i* of ***x*** with uniform probability and proposed a mutation of nucleotide *x_i_* to nucleotide 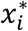 sampled from the set {*G*, *C*, *A*, *T*} – {*x_i_*} with uniform probability. The proposed mutation was accepted with probability given by the Metropolis criterion:

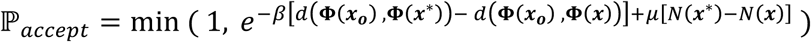

where ***x***^*∗*^ denotes the random variable ***x*** with the proposed mutation at index *i* [20].

To generate the results of Application 1, we used 50 chains in parallel for each ***x***_**0**_, with each chain constructed from 100 proposed mutations. Each chain was sampled after every proposal. To generate the results of Application 2, we used 100 Markov chains in parallel for each ***x***_**0**_, with each chain constructed from 3×10^4^ proposed mutations. Each chain was sampled every 100 proposals. To generate the results of Application 3, we used for each ***x***_**0**_ a single Markov chain constructed by proposing 1×10^6^ mutations. We sampled the chain every 100 proposals.

### Application 1: Interpreting learned XOR logic

#### Data set construction and network training

We trained instances of the architecture described in Application 1 on training sets of 2000 dinucleotides constructed by sampling i.i.d. multinomial distributions with P(A) = P(T) = 0.3 and P(G) = P(C) = 0.2, assuming independent bases. Training was performed using stochastic gradient descent with learning rate 5.0 ×10^−3^ and binary cross-entropy loss in the python package Keras [21]. After each training epoch, we evaluated the model on the set of 16 distinct dinucleotide inputs. Training was stopped when the model achieved 100% classification accuracy or after 40 epochs of training. Using this method, we trained 300 models and selected at random 30 of the 46 models achieving 100% accuracy. To insure stability of the learned solution (stochastic gradient descent of the loss does not guarantee non-decreasing classification accuracy), we trained the selected models for 4 additional epochs checking for 100% validation accuracy at each epoch and then applied the interpretation methods.

#### Network-dependent selection of *β*

We performed MaxEnt interpretation for all inputs to a single network using the same value of *β*. We selected *β* for each model by requiring that the width of MaxEnt samples be small relative to the empirical distribution of all dinucleotide inputs in the space of penultimate activations scaled by weights of connection to the output unit. Specifically, for each input ***x_i_*** to a fixed network, we solved for *β_i_* in the equation

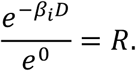

Here, *D* is a distance scale in the space of scaled penultimate activations at which the probability implied by distribution (2) has decayed by factor *R* and is defined as

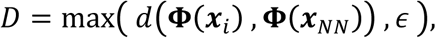

where *d* is the distance metric defined in (3) for a network with a single output unit and where ***Φ x***_*NN*_ denotes the nearest neighbor vector of penultimate activations in the empirical distribution of penultimate activations, and **ϵ** is a small value (on the scale of the empirical distance distribution measured by *d*) that handles the case of multiple inputs mapping to the same penultimate vector. Results in Application 1 use *R=*0.4 and **ϵ** = 1.0. We interpreted each network using *β* equal to the mean of all *β_i_*’s associated with the 16 dinucleotide inputs.

### Application 2: Localizing learned motifs

#### Data accession and network training

We downloaded the CTCF motif discovery data set from http://cnn.csail.mit.edu/motif_discovery/wgEncodeAwgTfbsHaibSknshraCtcfV0416102UniPk/ [2]. This data set was derived from a CTCF ChIP-seq experiment performed in human neuroblastoma cells treated with retinoic acid. Downloaded data were pre-partitioned into balanced training and test sets with 62,632 and 15,674 elements, respectively. We used this partition as a single fold in 5-fold cross validation (CV) scheme. For each CV fold, we set aside 1/8^th^ of training data for validation and trained the network architecture for 20 epochs with a batch size of 80, employing the categorical cross-entropy loss function and the adaDelta optimizer from python package Keras [21]. During training, but not during testing or interpretation, the dropout method with probability 0.1 was applied to the output of the fully connected layer to reduce overfitting [22]. We evaluated validation performance at the end of each epoch and selected the model with best validation accuracy.

#### Tomtom database query

We extracted MaxEnt motifs from the single nucleotide frequencies in Figure 3 A,B as the 19 bp intervals with the largest average KL divergence between sample nucleotide frequencies and a uniform distribution over A,C,G,T. The single nucleotide frequencies of the resulting motifs were then analyzed using the Tomtom webserver (version 4.12.0) [15] with default parameters.

#### MEME motif discovery and FIMO motif scan

We used the program MEME, version 4.10, to discover a consensus motif in 1500 inputs chosen randomly from the set of inputs where we applied our interpretation method [16]. We instructed MEME to find zero or one motif in each input with minimum and maximum motif lengths of 6 and 19, respectively. We required the consensus motif to be present in at least 400 inputs and stopped the search when the E-value exceeded 0.01.

We used the program FIMO, version 4.10, to scan the full set of 2500 sequences where we applied our interpretation [17]. We instructed FIMO not to search reverse complement strands of input sequences, and FIMO identified motifs in 1652 inputs. To construct the plot of relative distance distributions in Fig. 3 (C) and (D), whenever FIMO called more than one motif in a sequence, we measured the distance between the network-derived motif and the FIMO motif with the lowest p-value.

### Application 3: Extracting nucleosome positioning signals

#### Data set construction

We downloaded chemical cleavage maps of redundant and unique nucleosome dyads from the supplementary material of [10], and we used these dyad indices to construct data sets from the UCSC SAC2 version of the *S. cerevisiae* genome. Our positive validation and test data sets consisted of genomic intervals centered on unique nucleosome dyads of chromosomes 7 and 12, respectively. The positive training set consisted of genomic intervals centered on the set of redundant dyads from all other chromosomes (note that the set of redundant dyads contains the set of unique dyads). Each data set was balanced by negative elements centered on genomic indices sampled uniformly and without replacement from genomic indices at least 3 bps from all redundant dyads. This chromosome-based division of training, validation, and test data corresponded to roughly an 80%, 10%, 10% split of available data, and our test set contained 11738 elements. We represented all sequence inputs as one-hot arrays and then mean centered and standardized each entry by subtracting the genome-wide frequency *f* of the corresponding nucleotide and then dividing by 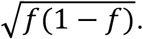 We found that this preprocessing gives a marginal improvement over simple one-hot encoding for this classification task.

#### Network training

We trained the network architecture using stochastic gradient descent with batch size of 8 and the categorical cross entropy loss function. Our training set was augmented with reverse complement sequences, and gradient descent used a learning rate of 0.1, momentum parameter of 0.5, and L2 weight penalty of 0.001. Training was done with the python package Keras [21].

#### Calculation of normalized Fourier amplitudes

Normalized Fourier amplitudes were calculated by performing discrete Fourier transform with the python package numpy [23], setting the zero frequency component to 0, then normalizing by the Euclidean norm of the Fourier components and calculating the amplitude at each frequency. These normalized amplitudes were averaged to produce the plots in Fig. 4(D).

#### Generating random orthogonal vectors for nearest neighbor classifier

We generated sets of 5 orthogonal basis vectors over the unit sphere embedded in 201 dimensions by sampling the 201 components of each vector from standard normal distributions and then performing QR decomposition on the 201 × 5 matrix of column vectors.

## Source Code and Data Availability

The source code, simulation data for Application 1, and a Python example workbook are available at https://github.com/jssong-lab/maxEnt.

## Acknowledgements and Funding

We thank Miroslav Hejna, Hu Jin and Wooyoung Moon for helpful discussions. This work has been supported by the National Science Foundation (DBI-1442504), the National Institutes of Health (R01CA163336), and the Founder Professorship from the Grainger Engineering Breakthroughs Initiative.

## Supporting information

**S1 Text. Details of Neural Network Layers**

**S2 Text. Derivation of MaxEnt distribution and input-wide feature importance score**

**S3 Text. Application of Saliency Map and DeepLIFT to network interpretation.**

**Figure S1 Interpretation of dinucleotide inputs to XOR network**

**Figure S2 Top hits for Tomtom database query with MaxEnt motifs**

**Figure S3 Saliency Map and DeepLIFT interpretation of CTCF bound sequences**

**Figure S4 Additional plots of nucleosome single nucleotide frequencies**

**Figure S5 Examples of learned low variance nucleosome features**

## References

1. Alipanahi B, Delong A, Weirauch MT, Frey BJ. Predicting the sequence specificities of DNA- and RNA-binding proteins by deep learning. Nat Biotechnol. 2015;33(8):831–8. doi: 10.1038/nbt.3300. PubMed PMID: 26213851.

2. Zeng H, Edwards MD, Liu G, Gifford DK. Convolutional neural network architectures for predicting DNA-protein binding. Bioinformatics. 2016;32(12):i121–i7. doi: 10.1093/bioinformatics/btw255. PubMed PMID: 27307608; PubMed Central PMCID: PMCPMC4908339.

3. Zhou J, Troyanskaya OG. Predicting effects of noncoding variants with deep learning-based sequence model. Nat Methods. 2015;12(10):931–4. doi: 10.1038/nmeth.3547. PubMed PMID: 26301843; PubMed Central PMCID: PMCPMC4768299.

4. Kelley DR, Snoek J, Rinn JL. Basset: learning the regulatory code of the accessible genome with deep convolutional neural networks. Genome Res. 2016;26(7):990–9. doi: 10.1101/gr.200535.115. PubMed PMID: 27197224; PubMed Central PMCID: PMCPMC4937568.

5. Zeng H, Gifford DK. Predicting the impact of non-coding variants on DNA methylation. Nucleic acids research. 2017. doi: 10.1093/nar/gkx177. PubMed PMID: 28334830.

6. Lanchantin J, Singh R, Wang B, Qi Y. Deep Motif Dashboard: Visualizing and Understanding Genomic Sequences Using Deep Neural Networks. Pac Symp Biocomput. 2016;22:254–65. PubMed PMID: 27896980.

7. Simonyan K, Vedaldi A, Zisserman A, editors. Deep Inside Convolutional Networks: Visualising Image Classification Models and Saliency Maps. ICLR Workshop 2014.

8. Shrikumar A, Greenside P, Shcherbina A, Kundaje A. Not Just a Black Box: Learning Important Features Through Propagating Activation Differences. 2016;(arXiv:1605.01713 [cs.LG]).

9. Lundberg S, Lee S-I. An unexpected unity among methods for interpreting model predictions. NIPS 2016 Workshop on Interpretable Machine Learning in Complex Systems 2016.

10. Brogaard K, Xi L, Wang JP, Widom J. A map of nucleosome positions in yeast at base-pair resolution. Nature. 2012;486(7404):496–501. Epub 2012/06/23. doi: 10.1038/nature11142. PubMed PMID: 22722846; PubMed Central PMCID: PMCPMC3786739.

11. Consortium EP. An integrated encyclopedia of DNA elements in the human genome. Nature. 2012;489(7414):57–74. doi: 10.1038/nature11247. PubMed PMID: 22955616; PubMed Central PMCID: PMCPMC3439153.

12. Ohlsson R, Renkawitz R, Lobanenkov V. CTCF is a uniquely versatile transcription regulator linked to epigenetics and disease. Trends Genet. 2001;17(9):520–7. Epub 2001/08/30. PubMed PMID: 11525835.

13. Chan CS, Song JS. CCCTC-binding factor confines the distal action of estrogen receptor. Cancer Res. 2008;68(21):9041–9. Epub 2008/11/01. doi: 10.1158/0008-5472.CAN-08-2632. PubMed PMID: 18974150.

14. Crooks GE, Hon G, Chandonia JM, Brenner SE. WebLogo: a sequence logo generator. Genome Res. 2004;14(6):1188–90. doi: 10.1101/gr.849004. PubMed PMID: 15173120; PubMed Central PMCID: PMCPMC419797.

15. Gupta S, Stamatoyannopoulos JA, Bailey TL, Noble WS. Quantifying similarity between motifs. Genome Biol. 2007;8(2). doi: ARTN R24 10.1186/gb-2007-8-2-r24. PubMed PMID: WOS:000246076300019.

16. Bailey TL, Johnson J, Grant CE, Noble WS. The MEME Suite. Nucleic acids research. 2015;43(W1):W39–49. Epub 2015/05/09. doi: 10.1093/nar/gkv416. PubMed PMID: 25953851; PubMed Central PMCID: PMCPMC4489269.

17. Grant CE, Bailey TL, Noble WS. FIMO: scanning for occurrences of a given motif. Bioinformatics (Oxford, England). 2011;27(7):1017–8. Epub 2011/02/19. doi: 10.1093/bioinformatics/btr064. PubMed PMID: 21330290; PubMed Central PMCID: PMCPMC3065696.

18. Jin H, Rube HT, Song JS. Categorical spectral analysis of periodicity in nucleosomal DNA. Nucleic acids research. 2016;44(5):2047–57. doi: 10.1093/nar/gkw101. PubMed PMID: 26893354; PubMed Central PMCID: PMCPMC4797311.

19. Hughes AL, Rando OJ. Mechanisms underlying nucleosome positioning in vivo. Annu Rev Biophys. 2014;43:41–63. doi: 10.1146/annurev-biophys-051013-023114. PubMed PMID: 24702039.

20. Metropolis N, Rosenbluth AW, Rosenbluth MN, Teller AH, Teller E. Equation of State Calculations by Fast Computing Machines. J Chem Phys. 1953;21(6):1087–92.

21. Chollet F. Keras: GitHub; 2015. Available from: https://github.com/fchollet/keras.

22. Srivastava N, Hinston G, Krizhevasky A, Sutskever I, Salakhutdinov R. Dropout: A Simple Way to Prevent Neural Networks from Overfitting. Journal of Machine Learning Research 2014;15:1929–58.

23. van der Walt S, Colbert C, Varoquaux G. The NumPy array: a structure for efficient numerical computation. Computing in Science and Engineering. 2011;13(2):22–30.

